# Analysis of Biofilm Complexity in 3D (ABC3D): An open-source framework for quantitative fractal, textural, and statistical analysis of colony biofilm morphology in three dimensions

**DOI:** 10.64898/2026.02.27.708470

**Authors:** Gail McConnell

## Abstract

2.

Quantitative image analysis is central to understanding microbial growth, morphology, and spatial organisation. However, conventional metrics such as mean intensity or object count often do not capture the complex structural heterogeneity and patterning characteristic of microbial colonies and biofilms. To address this limitation, Analysis of Biofilm Complexity in 3D (ABC3D), an open-source Python framework for automated extraction of fractal, textural, and statistical descriptors from volumetric microscopy images, is reported. ABC3D computes a set of parameters including fractal dimension, lacunarity, entropy, grey-level co-occurrence matrix features, and wavelet sub-band energies from three-dimensional (3D) image datasets. ABC3D is demonstrated in macrocolony biofilms formed by cell shape mutants of *Escherichia coli*, where it is shown that nutrient availability accounts for the majority of structural variance, while cell shape produces additional structural variation that differs between nutrient conditions. ABC3D provides researchers with an accessible, quantitative approach to assessing biofilm morphology in microscopy datasets.

**Summary:** An open-source, quantitative analysis pipeline is presented that integrates fractal, lacunarity, entropy, texture and wavelet descriptors to characterise colony biofilm architecture in three dimensions. Application to *Escherichia coli* cell shape mutants demonstrates that macrocolony biofilm architecture is best understood as a coordinated, multiscale phenotype rather than as an aggregate of independent structural metrics.

**Impact statement:** Biofilm architecture is pivotal for microbial survival, antimicrobial tolerance, and ecological function but tools to quantify structural organisation in these cell communities remain limited. The commonest metrics describe bulk properties such as width, thickness, or cell number, but they do not capture multiscale spatial heterogeneity. Here, an open-source framework for Analysis of Biofilm Complexity in 3 Dimensions (ABC3D) is reported. This software integrates measurements of fractal geometry, lacunarity, entropy, texture statistics, and wavelet energy. ABC3D is demonstrated in *Escherichia coli* macrocolony biofilms, where it is shown that nutrient environment has a leading role in determining colony architecture in *E. coli* biofilms, while cell shape has a lesser but still significant influence on structural variation. The ABC3D pipeline can be applied to any microbial communities imaged by confocal microscopy and other volumetric imaging methods and has the potential to give a deeper understanding of how cells organise in biofilms.

**Data summary:** Full code for ABC3D and data analysis is available at https://github.com/gailmcconnell/ABC3D. Image data are available upon request.

The author confirms all supporting data, code and protocols have been provided within the article or through supplementary data files.

## 5. Introduction

Understanding how biofilms are structured is essential for interpreting how they function and respond to environmental stress. Confocal laser scanning microscopy (CLSM) has made high-resolution imaging of biofilms possible in two dimensions (2D) and three dimensions (3D) (1,2), but the field has been slow to develop the software tools necessary for measurement of these complex living structures. Early tools such as COMSTAT provided fundamental but important measurements from CLSM data such as biofilm volume and surface-to-volume ratio, and these enabled comparisons of biofilm architecture in different environmental conditions (3). COMSTAT, and subsequently COMSTAT2, (4) has helped to establish a solid basis for morphological measurements of biofilms.

Further image data approaches such as digital image in microbial ecology (daime) (5) and later Biofilm Intensity and Architecture Measurement (BIAM) (6) extended measurement capability to include segmentation, 2D/3D object quantification, and visualisation for microbial communities. More recently, BiofilmQ has extended capability further, offering cell segmentation and measurement in space and time (7).

Biofilm architecture maps directly onto mass-transport, reaction, and detachment phenomena (8,9), and thicker biofilms have been reported to harbour deeper gradients of oxygen and nutrients, which can modulate growth (10–12). Measurement of these properties has assisted in the understanding the relationship between architecture and function (13).

Architecture also shapes antimicrobial tolerance. The extracellular matrix (EPS) can impede diffusion and create chemical microenvironments that inhibit growth and change antibiotic susceptibility (14). Increases in thickness, roughness, and porosity often correlate with reduced penetration and elevated tolerance (13).

Beyond gross morphology, texture metrics, including fractal dimension, lacunarity, grey-level co-occurrence matrix (GLCM) statistics, entropy, and multi-scale spectral measures, capture heterogeneity and self-similarity across scales (15–18), and are distinct from volume and thickness. To aid interpretation of these descriptors, Figure 2 provides a conceptual illustration of how each parameter varies from low to high values in representative biofilm cross-sections. These schematics highlight the distinct structural properties captured by fractal, lacunarity, entropy, texture, and wavelet-based energy measures. Fractal dimension quantifies how completely a structure fills space across scales; lacunarity captures the distribution and size of gaps; entropy measures intensity heterogeneity; grey-level co-occurrence statistics describe local contrast and spatial correlation; and wavelet sub-band energy quantify how structural variation is distributed across spatial frequencies and directions. Together, these descriptors distinguish dense, homogeneous architectures from heterogeneous, channel-rich structures. Early work to extend fractal and texture-based analysis to 3D biofilm datasets included Image Structure Analyzer in 3D (ISA3D) software, which measured the box-counting fractal dimension, lacunarity, and grey-level texture of confocal image stacks (19). ISA3D demonstrated that volumetric biofilm complexity could be quantified beyond traditional bulk parameters and this represented an important methodological advance at the time. However, the software was distributed as MATLAB-based code and is no longer actively maintained or publicly accessible. Contemporary tools increasingly support these measurements, but workflows typically still require stitching together of multiple libraries or software apps. These limitations highlight the need for a transparent technology for multiscale three-dimensional architectural phenotyping.

Analysis of Biofilm Complexity in 3D (ABC3D) is introduced as an open-source Python framework that unifies a broad range of quantitative descriptors relevant to biofilm image data analysis. It consolidates fractal measures, textural features derived from grey-level co-occurrence matrices, energy parameters, and statistical analyses within a single workflow. By analysing 3D data, ABC3D overcomes limitations of two-dimensional (2D) analysis. This framework provides microbiologists with a flexible and extendable tool for linking biofilm architecture and texture to underlying physiological and ecological processes.

## 6. Materials and Methods

A schematic diagram showing the experimental and computational pipeline is shown in Figure 1.

**Figure 1.**
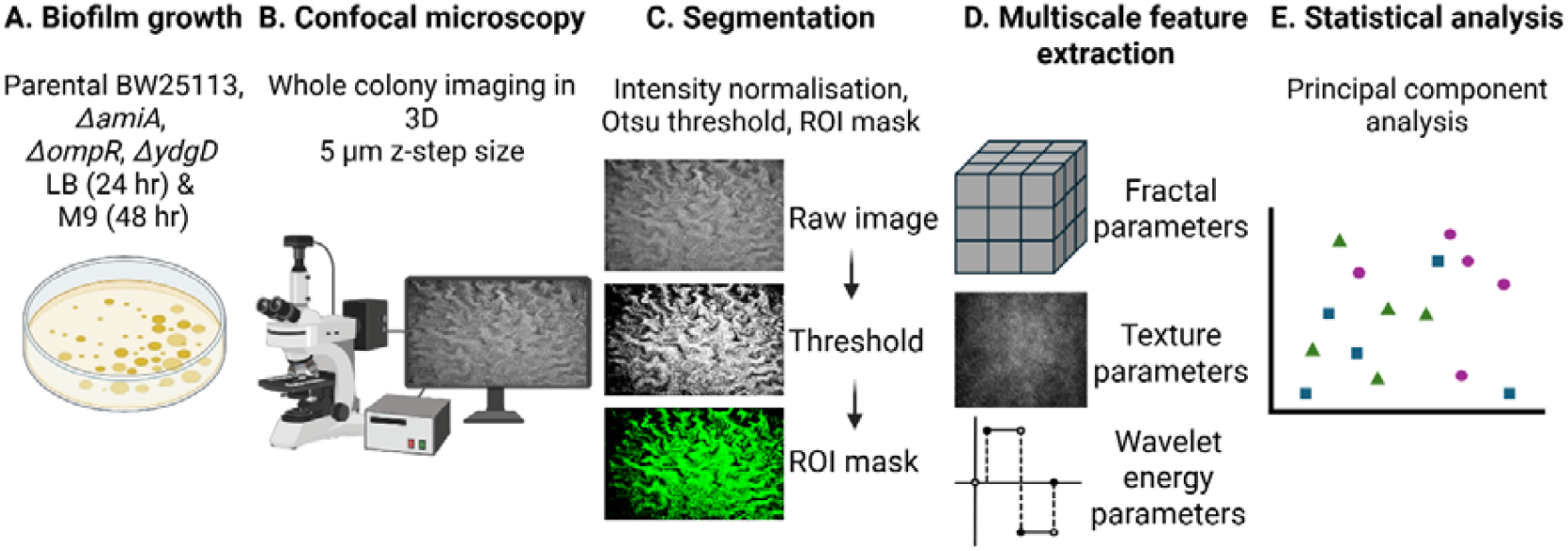
Experimental and computational workflow for multiscale biofilm architecture analysis. A. Biofilm growth conditions. Parental (BW25113) and deletion mutants (Δ*amiA*, Δ*ompR*, Δ*ydgD*) were grown as single-founder, isogenic microcolony biofilms on LB agar (24 hr) and M9 minimal agar (48 hr). B. Whole colony three-dimensional image stacks were acquired by confocal laser scanning microscopy using a 5 μm z-step size to capture full colony architecture. C. Raw volumetric images were intensity normalised and segmented using Otsu thresholding to generate a binary region-of-interest (ROI) mask representing biofilm biomass. D. Quantitative descriptors were computed from segmented volumes, including fractal parameters, texture features derived from grey-level co-occurrence matrices, and wavelet sub-band energy metrics. E. Feature matrices were standardised, and principal component analysis (PCA) was performed to identify dominant axes of structural variation and quantify the effects of growth medium and strain on biofilm architecture. Created with BioRender.com.

**Figure 2.**
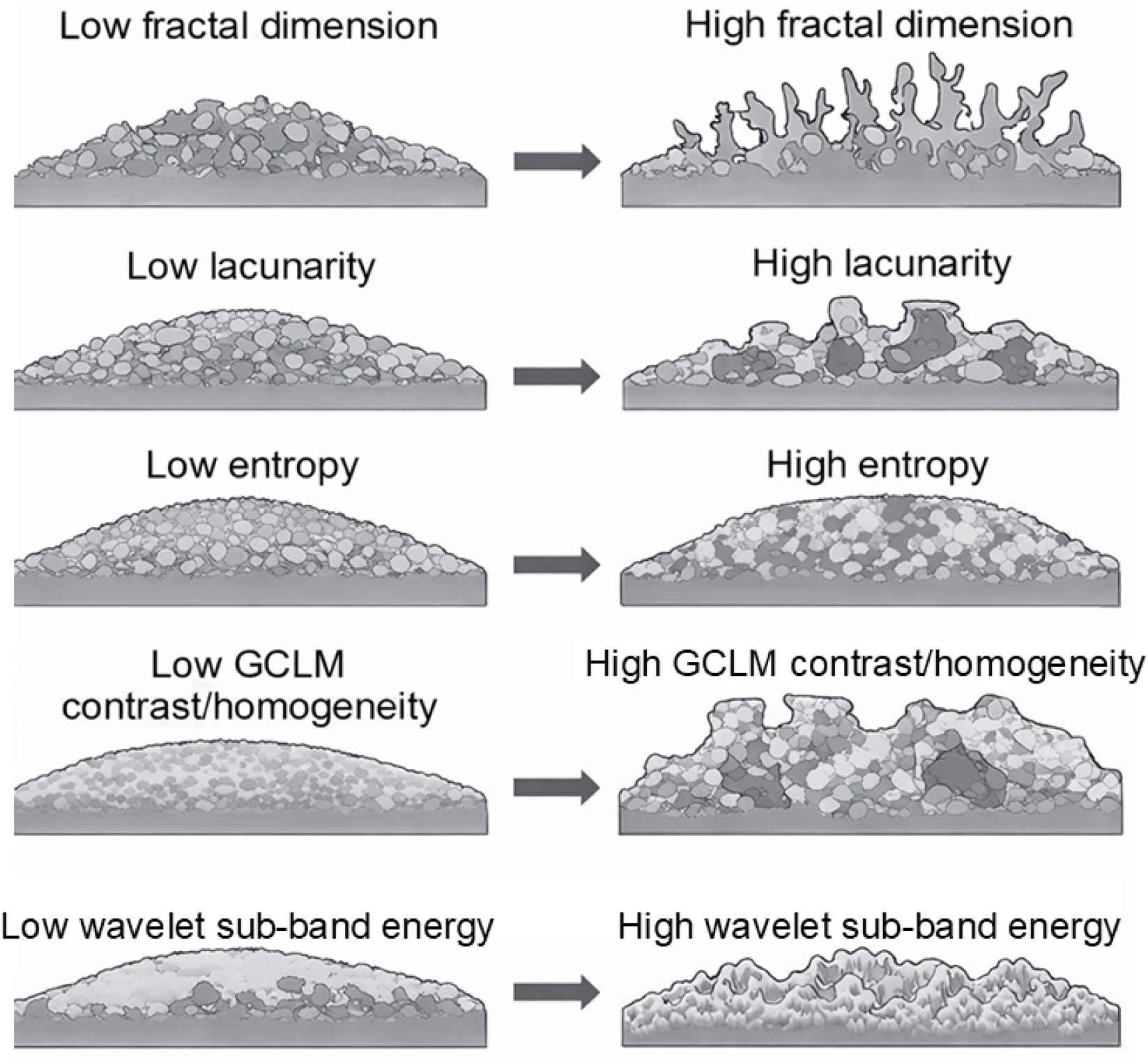
Schematic illustration of multiscale architectural descriptors computed by ABC3D. Schematic cross-sectional (xz) representations of biofilm structure demonstrating how key quantitative descriptors vary from low (left) to high (right) values. Increasing fractal dimension reflects greater scale-invariant structural complexity and space-filling branching. Increasing lacunarity indicates greater heterogeneity in gap size and spatial distribution of voids. Increasing entropy represents greater voxel-level intensity heterogeneity within the biofilm volume. Increasing GLCM contrast reflects stronger local intensity differences between neighbouring voxels and greater textural roughness. Increasing wavelet sub-band energy represents greater magnitude of structural variation at defined spatial frequencies, corresponding to fine-scale irregularities. Illustrations are conceptual and intended to provide intuitive interpretation of quantitative metrics rather than report direct biological measurements.

### 6.1. Software architecture and computation of features

ABC3D (Analysis of Biofilm Complexity in 3D) is a standalone, script-based workflow written in Python (v3.11.13). The analysis was implemented as custom Python code using libraries from the scientific Python ecosystem, including NumPy, pandas, tifffile, scikit-image, SciPy, and PyWavelets. Code development and execution were performed in the Spyder (v6.0.8) integrated development environment and the software was run on a HP EliteBook 830 laptop computer with an Intel® Core™ Ultra 5 1.6 MHz processor and 16 GB RAM, with a Microsoft Windows 11 Enterprise operating system.

Three-dimensional OME-TIFF stacks are imported and standardised to (z, y, x) ordering. Intensity data are converted to 32-bit floating-point format to ensure numerical stability and memory efficiency. Biofilm biomass is segmented from background using global intensity thresholding. Otsu’s method is applied to a reproducible random subsample of up to 10□ voxels rather than the full volume (20). This sampling limit is set to control processing time, rather than being a memory constraint. Optional parameters allow manual threshold specification, and removal of small, connected components below a user-defined voxel size to suppress noise. The resulting binary mask defines the biofilm region of interest (ROI) for all subsequent analyses.

Structural quantification was performed directly on the segmented 3D volume. Total foreground voxel count and foreground fraction were calculated to estimate biomass occupancy within the imaged volume. Intensity heterogeneity within the volume was quantified using Shannon entropy and Rényi entropy (α = 2) (15,21).

Scale-dependent architectural complexity was assessed using fully three-dimensional box-counting analysis and lacunarity. Fractal dimension was obtained via Relative Dimension Box-Counting (RDBC) from log–log regression across dyadic box sizes, with R^2^ reported as a goodness-of-fit metric (22). Lacunarity was computed across equivalent spatial scales to characterise structural heterogeneity and gap distribution within the biofilm (17).

Local spatial organisation was evaluated using slice-wise masked grey-level co-occurrence matrices (2.5D GLCM), from which Haralick statistics (angular second moment, contrast, correlation, dissimilarity, energy, and homogeneity) were summarised across slices (16).

Multiscale structural organisation was further characterised using a single-level three-dimensional discrete wavelet transform (Daubechies db2). Energy was computed for each 3D sub-band (LLL, LLH, LHL, LHH, HLL, HLH, HHL, HHH), where H=high and L=low, providing directional multiscale measurements of architectural variation (18,23,24).

All computed features were exported as a single.CSV file. All analysis parameters were explicitly defined within the script to ensure full reproducibility. Full mathematical definitions of all computed descriptors are provided in Supplementary Information.

### 6.2. Experimental design and image data acquisition

No new image data were acquired for demonstrating ABC3D. Instead, data were repurposed from Bottura *et al*, where full information on specimen preparation is available (25). A brief overview of specimen preparation and image data acquisition is presented here.

The bacterial strains used were obtained from the Keio collection (26), a single-gene knockout library of all nonessential genes in the *E. coli* K-12 strain, BW25113 (27). Green fluorescent protein (GFP)-expressing strains parental BW25113/pAJR145, BW25113 Δ*amiA*::kan/pAJR145, BW25113 Δ*ompR*::kan/pAJR145 and BW25113 Δ*ydgD*::kan/pAJR145 referred to in the text as Δ*amiA*, Δ*ompR*, and Δ*ydgD*, were selected for their modified cell phenotype. Macrocolony biofilms of each strain were grown on agar substrates in a sterile 3D-printed plastic specimen holder, as described previously (28). Macrocolony biofilms were grown for 24□h when using Miller’s lysogeny broth (LB) and for 48□h when using M9 minimal medium supplemented with 0.2% glucose. This difference in growth is due to the doubling time of *E. coli* in M9 being half that of the doubling time in LB (29).

Mature colony biofilm images were acquired on an Olympus IX81 microscope coupled to a FluoView FV1000 confocal laser scanning unit (Olympus, Japan). Fluorescence from GFP was excited using a 488 nm argon laser (GLG3135, Showa Optronics, Japan) and was detected by a photomultiplier tube (PMT) with a spectral detection window set between wavelengths of 510 and 560□nm. Samples were imaged using a 10×/0.4 dry objective lens for resolving sub-cellular structure.

Three-dimensional z-stacks of colony biofilms grown on LB and M9 media were acquired with a slice spacing of 5□µm for Nyquist sampling in the axial dimension. Colony biofilms were imaged for each strain and each medium (n=5 repeats). All data were saved in the proprietary.oib file format and were converted to OME.TIFF in FIJI (30). Representative maximum intensity projection images, also made in FIJI, of each strain and nutrient are shown in Figure 3.

**Figure 3.**
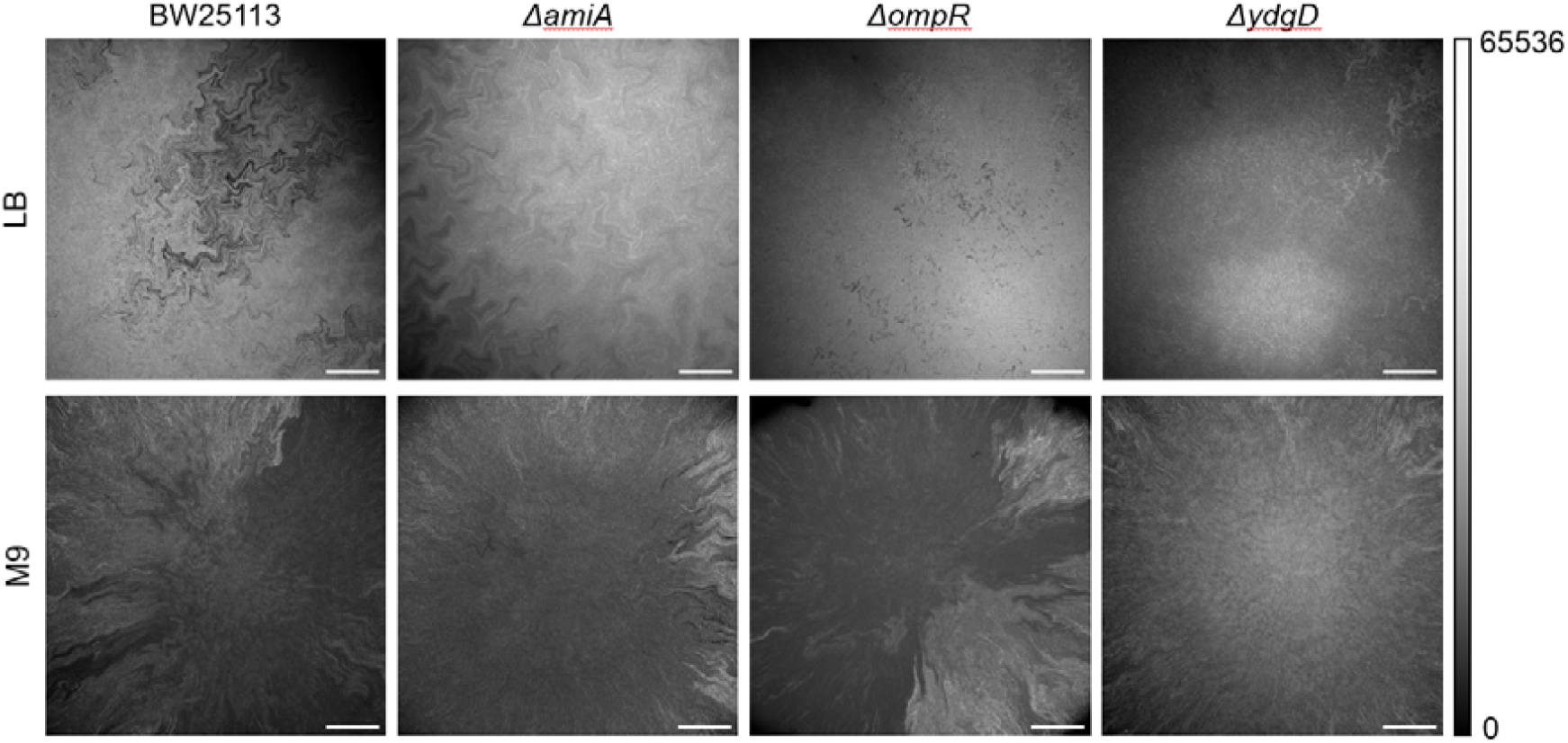
Representative maximum intensity projection images of confocal microscopy datasets showing the morphology of biofilms formed by the strains BW25113 (parental), Δ*amiA*, Δ*ompR*, and Δ*ydgD* in LB and M9 media. Scale bar = 200 µm.

### 6.3. Statistical analysis

All statistical analyses of all image datasets were performed in Python using scikit-learn (31) and statsmodels (32). Quantitative features derived from ABC3D (entropy measures, fractal dimension, lacunarity, GLCM statistics, and wavelet sub-band energies) were assembled into a single dataset data matrix in which each row represented one imaged biofilm, and each column represented one measured parameter, e.g. lacunarity, Shannon entropy etc. This matrix formed the basis for multivariate statistical analysis.

#### 6.3.1. Principal component analysis

Principal component analysis (PCA) was used to simplify the dataset by summarising multiple architectural measurements into a smaller number of composite variables (33). Before PCA, all features were standardised using zero-mean, unit-variance scaling so that measurements with larger numerical ranges did not disproportionately influence the analysis.

PCA transforms the original correlated features into new, independent variables called principal components (PCs). Each principal component represents a distinct pattern of variation in biofilm architecture. For each biofilm, a PC score indicates its position along a given component axis, effectively summarising how strongly that biofilm expresses the structural behaviour described by that component.

Each component explains a proportion of the total variance in the dataset. Components are ordered from highest to lowest explained variance, meaning that the first component (PC1) captures the largest share of overall architectural variation across biofilms, the second component (PC2) captures the next largest share, and so on. The first two components (PC1 and PC2) were retained for visualisation and subsequent statistical analysis, as together they accounted for most of the total variance.

Feature loadings were examined to identify which architectural measurements contributed most strongly to separation along each component axis.

#### 6.3.2. Two-way ANOVA on Principal Components

To assess the contributions of genetic background and growth medium to biofilm architecture, two-way ANOVA models were fitted to PC1 and PC2 scores using ordinary least squares regression with strain and medium as fixed factors, including a strain × medium interaction term. Type II sums of squares were used to evaluate main effects in the presence of interaction. All tests were two-sided, and statistical significance was defined as P < 0.05.

#### 6.3.3. Per-feature ANOVA and Multiple Testing Correction

The same two-way ANOVA framework was applied independently to each architectural feature. For each model, F-statistics and corresponding P-values were obtained. To account for multiple testing across features, false discovery rate (FDR) correction was performed using the Benjamini-Hochberg procedure (34), applied separately to medium, strain, and interaction terms. Adjusted q-values < 0.05 were considered significant.

#### 6.3.4. Data presentation

PCA scatter plots, interaction plots (showing group means ± standard error of the mean, SEM), and heatmaps of mean z-scored feature values grouped by strain × medium were generated from ABC3D outputs using a custom Python code.

ANOVA results are reported as *F*(*df*_l_, *df*_2_) = value, P = value, where *df*_l_ and *df*_2_ denote numerator and denominator degrees of freedom, respectively. All statistical tests were two-sided unless otherwise stated.

## 7. Results

### 7.1. Multiscale feature extraction with ABC3D captures dominant axes of architectural variation

Biofilm architecture was measured across genetic backgrounds and nutrient conditions by extracting structural features from 3D confocal stacks using ABC3D.

Principal component analysis (PCA) of the full parameter set revealed that PC1 related to 46.3% of total variance and PC2 related to 26.8% of the total variance, together accounting for 73.1% of architectural variability across all samples. This is shown in Figure 4. PC3 accounted for 8.4% of total variance and did not materially alter clustering patterns. As such, visualisation and interpretation focused on PC1-PC2 space. This result indicates that most of the structural heterogeneity can be represented in a 2D principal component space derived from the full multidimensional image data.

**Figure 4.**
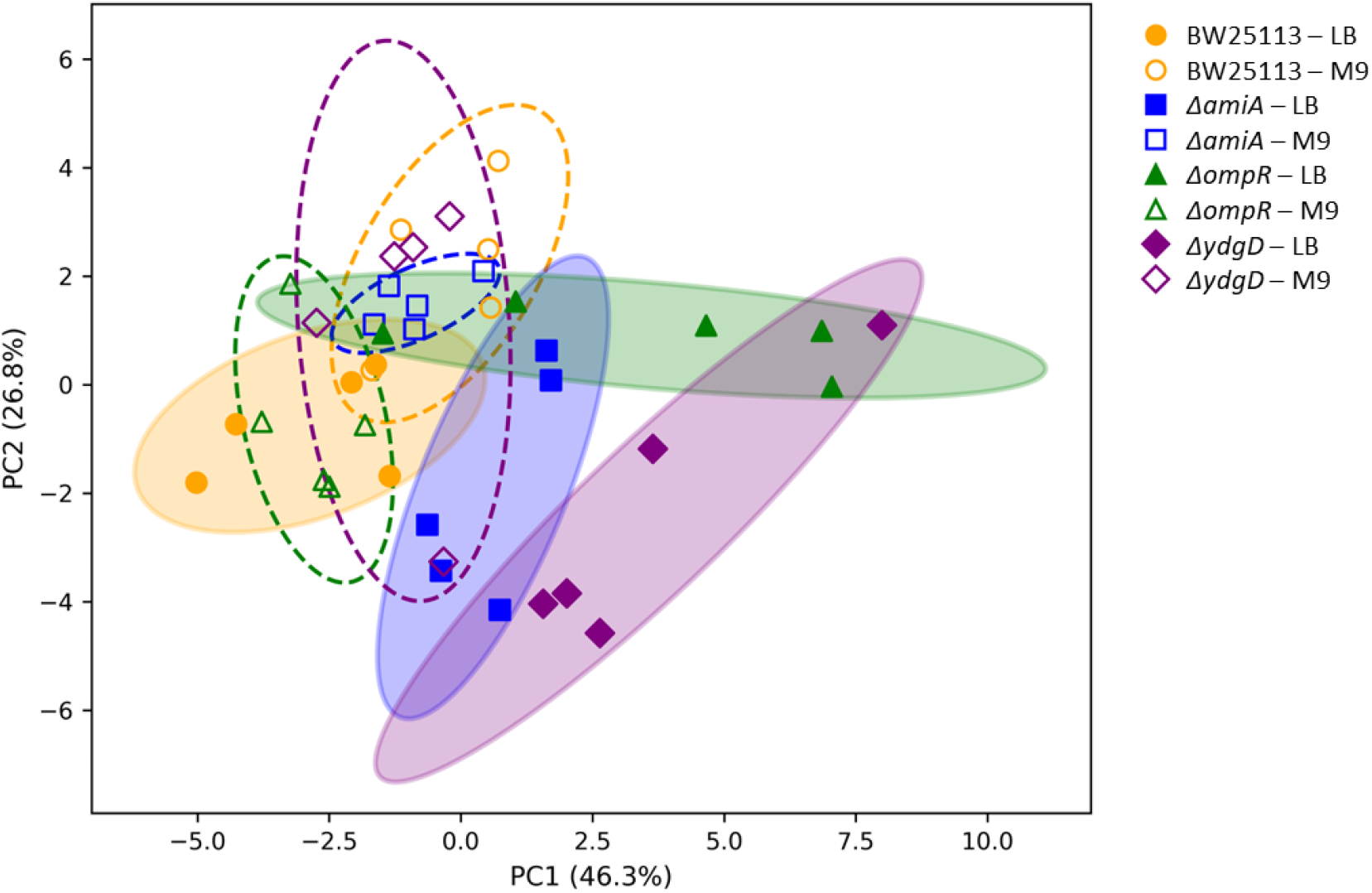
Principal component analysis (PCA) of standardised full-feature biofilm descriptors derived from n=5 3D confocal image stacks using ABC3D. PC1 accounts for 46.3% of total variance and PC2 accounts for 26.8%. Separation along PC1 primarily distinguishes strains, whereas PC2 further resolves medium-dependent architectural responses. Ellipses represent covariance-based data ellipses (2 standard deviations) calculated from the PC1 and PC2 scores for each strain × medium group, illustrating the within-group dispersion in PCA space.

### 7.2. Medium and cell shape jointly influence biofilm architecture

Projection of samples into PC space revealed structured separation by both growth medium and cell shape (Figure 4). Mean PC1 values (± standard deviation, SD) for LB-grown colonies were: parental BW25113 = 2.87 ± 1.66, Δ*amiA* = 0.61 ± 1.10, Δ*ompR* = 3.62 ± 3.74, and Δ*ydgD* = 3.57 ± 2.59. In M9, corresponding PC1 means were parental BW25113 = −0.21 ± 1.12, Δ*amiA* = −0.86 ± 0.80, Δ*ompR* = −2.79 ± 0.75, and Δ*ydgD* = −1.09 ± 1.02. These values indicate that PC1 shifts directionally between nutrient conditions for all strains, with the largest magnitude change observed for Δ*ompR*. Notably, ΔompR colonies grown in LB exhibited greater dispersion along PC1 (SD = 3.74) compared with other groups, indicating increased within-condition architectural variability.

PC2 further resolved cell shape and medium-dependent differences. For LB-grown colonies, mean PC2 values (± SD) were: parental BW25113 = 0.76 ± 0.98, Δ*amiA* = 1.89 ± 2.13, Δ*ompR* = −0.90 ± 0.58, and Δ*ydgD* = 2.51 ± 2.41. In M9, PC2 means were: parental BW25113 = −2.23 ± 1.46, Δ*amiA* = −1.50 ± 0.46, Δ*ompR* = 0.64 ± 1.50, and Δ*ydgD* = −1.17 ± 2.58. These shifts demonstrate that cell shape dependent architectural responses are modulated by nutrient environment rather than fixed across conditions.

To formally assess contributions of medium and cell shape, a two-way ANOVA was performed on PC scores. For PC1, there was a significant dependence on nutrient availability (*F*_l,32_ = 17.30, P < 0.001), a significant dependence on strain (*F*_3,32_ = 3.87, P = 0.018), and a significant medium × strain interaction (*F*_3,32_ = 11.26, P < 0.001). The large interaction term indicated that macrocolony architecture depended most strongly on nutrient availability.

For PC2, there was a significant dependence on nutrient availability medium (*F*_l,32_ = 15.83, P< 0.001), no significant dependence on strain (*F*_3,32_ = 1.21, P = 0.321), and a significant medium × strain interaction (*F*_3,32_ = 5.31, P = 0.004). As such, while the growth medium alone strongly influenced PC2, strain-specific effects emerged primarily through interaction with medium.

Interaction plots shown in Figure 5 illustrate these cross-over effects. For PC1, Δ*ompR* and Δ*ydgD* decreased sharply from LB to M9, whereas WT increased and Δ*amiA* showed a decrease. The crossing trajectories confirmed non-additive cell shape-environment effects. For PC2, parental BW25113, Δ*amiA*, and Δ*ydgD* increased in LB relative to M9, whereas Δ*ompR* decreased in LB relative to M9, again demonstrating opposing conditional architectural responses.

**Figure 5.**
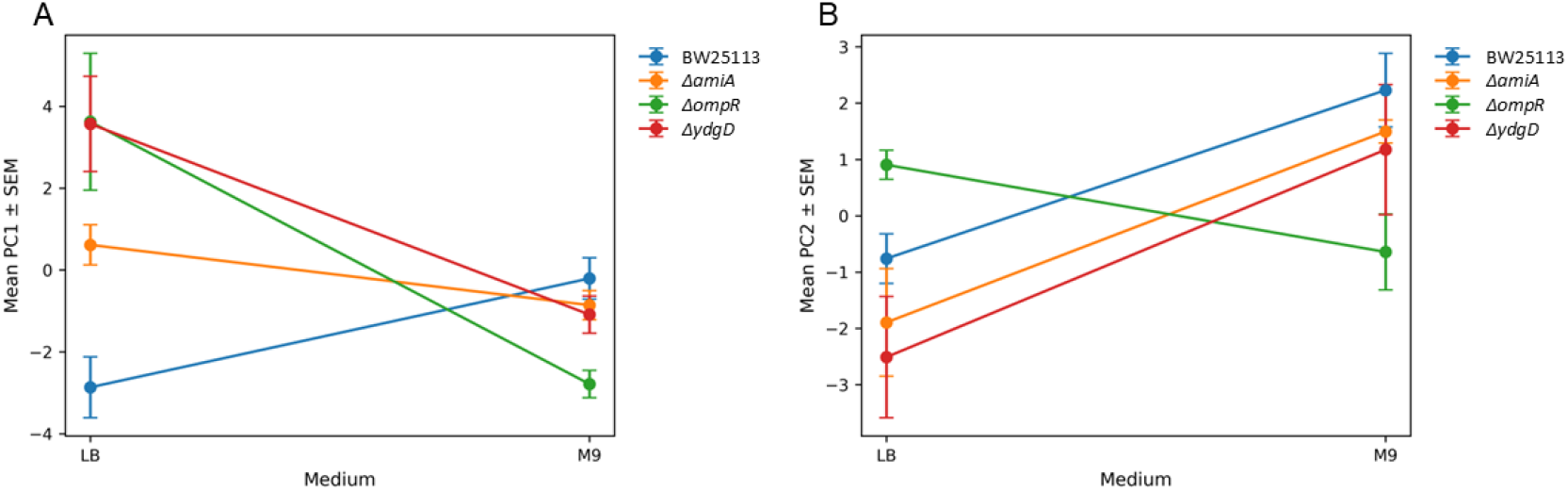
Strain × medium interaction effects on principal component scores. A. PC1 interaction plot showing mean PC1 scores (± SEM) for each strain grown in LB or M9. PC1 captures 46.3% of total variance and reflects large-scale architectural organisation dominated by wavelet sub-band energy and textural homogeneity features. Parental BW25113 shifts from negative PC1 in LB to near-zero in M9, whereas Δ*ompR* and Δ*ydgD* display strong reductions from LB to M9. Δ*amiA* shows a modest decrease across media. Divergent slopes indicate a strain × medium interaction effect on the dominant structural axis. B. PC2 interaction plot. Mean PC2 scores (± SEM) for each strain and medium. PC2 accounts for 26.8% of variance and reflects complementary texture-heterogeneity trade-offs. Parental BW25113, Δ*amiA*, and Δ*ydgD* shift positively from LB to M9, whereas Δ*ompR* decreases. The opposing directional response of Δ*ompR* relative to other strains indicates a distinct environmental adaptation phenotype.

These results established that biofilm structure is governed by strong cell shape-environment interactions.

### 7.3. PC1 reports multiscale energy concentration and structural homogeneity

The fifteen largest absolute loadings are shown in Figure 6. PC1 was dominated by positive contributions from multiple wavelet sub-band energy features (LLL, HHH, LHH, LLH), together with GLCM energy, angular second moment, and homogeneity, as well as RDBC. Negative loadings were primarily associated with lacunarity and GLCM contrast-based measures (correlation, dissimilarity, and contrast). These results indicated that PC1 represented a continuum between high PC1 values, namely high multiscale wavelet sub-band energy, high GLCM energy and homogeneity, higher fractal dimension, and lower lacunarity and texture contrast, as expected in dense, energetically concentrated structures.

**Figure 6.**
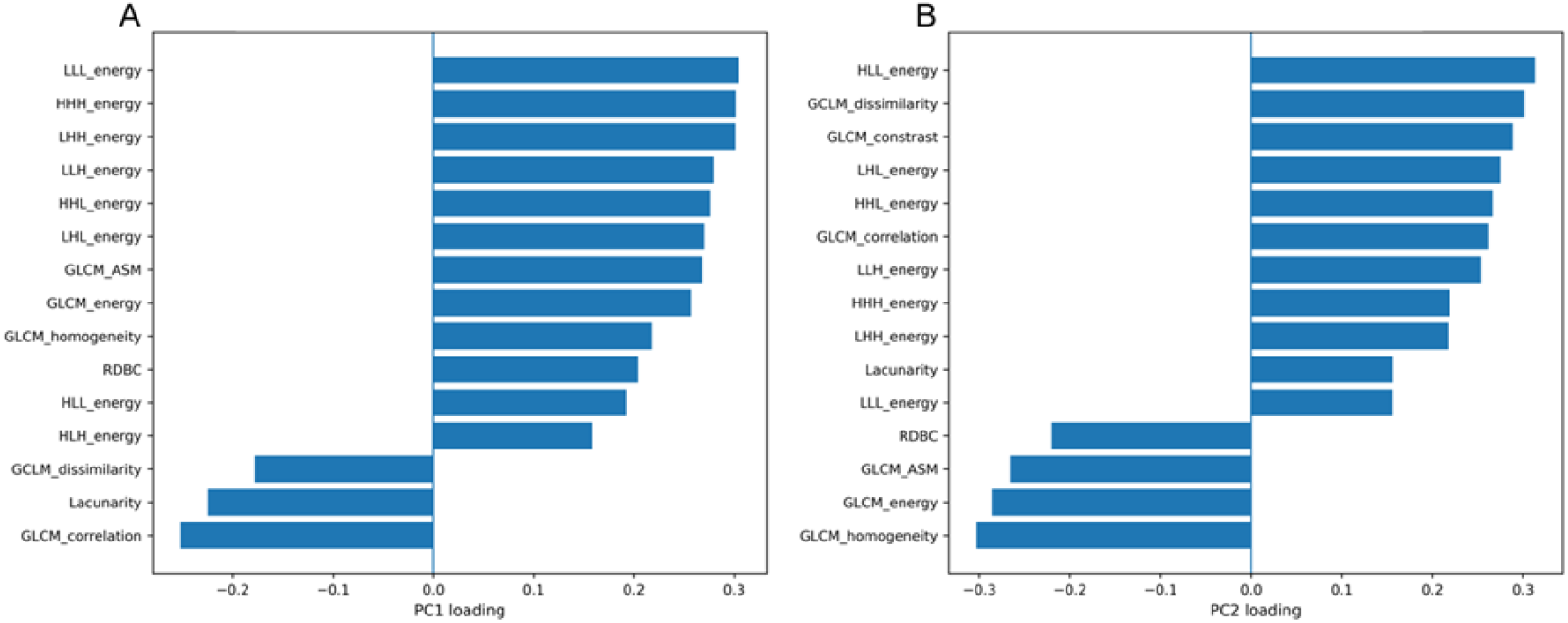
Principal component loadings reveal multiscale structural determinants of biofilm variation. A. PC1 is dominated by positive contributions from multiscale wavelet sub-band energy features (LLL energy, HHH energy, LHH energy, LLH energy, HHL energy, LHL energy), alongside GLCM angular second momentum and GLCM energy. Negative loadings are observed for GLCM correlation, lacunarity, and GLCM dissimilarity. PC1 therefore represents a gradient from correlated, heterogeneous structures to energetically dominant, spatially organized architectures. B. PC2 loadings indicate strong positive contributions from HLL energy, GLCM dissimilarity, GLCM contrast, and GLCM correlation, while strong negative loadings arise from GLCM homogeneity, GLCM energy, and GLCM angular second momentum. This axis reflects a contrast-homogeneity trade-off and captures medium-dependent reorganisation of texture features.

Low PC1 values, including higher lacunarity, greater pixel-to-pixel contrast and dissimilarity, and stronger correlation structure were consistent with more spatially heterogeneous architectures. Δ*ompR* and Δ*ydgD* colonies grown on LB occupy the high-PC1 end of this spectrum, corresponding to dense and relatively homogeneous structures, whereas parental BW25113 in LB lies toward the lower-PC1 end, reflecting greater heterogeneity. In M9, this pattern is reversed, indicating that architecture depends most strongly on nutrient availability.

### 7.4. PC2 captures intermediate-scale structure and contrast

The largest PC2 loadings defined a second, independent axis of structural variation. Positive PC2 loadings were dominated by intermediate-frequency wavelet sub-band energies (HLL, LHL, HHL) together with GLCM contrast and dissimilarity. Negative loadings were associated with GLCM homogeneity, angular second moment, energy, and RDBC.

PC2 therefore distinguishes architectures characterised by increased intermediate-scale structural variation and local contrast from those with greater homogeneity and energy distribution.

On M9, parental BW25113, Δ*amiA*, and Δ*ydgD* shift toward higher PC2 values, indicating increased heterogeneity under nutrient limitation. In contrast, Δ*ompR* shifts negatively along PC2 in M9, suggesting that it has a more homogeneous structural state. This divergence indicates that strains do not respond uniformly to nutrient limitation.

### 7.5. Multiscale parameters shift across cell shape and environment

The mean z-score heatmap by strain × medium (Figure 7) indicates coordinated, scale-dependent organisation. On LB, Δ*ompR* showed broadly elevated wavelet sub-band energy features and increased entropy measures, while Δ*ydgD* exhibited elevated GLCM angular second momentum and GLCM energy with reduced entropy and contrast, which is consistent with increased local ordering. On M9, parental BW25113 biofilms displayed increased wavelet sub-band energy with moderate homogeneity. In contrast, Δ*ompR* showed reduced wavelet sub-band energy and Shannon entropy with a shift toward greater homogeneity, whereas Δ*ydgD* exhibited elevated Shannon entropy and GLCM contrast relative to LB, indicating increased heterogeneity.

**Figure 7.**
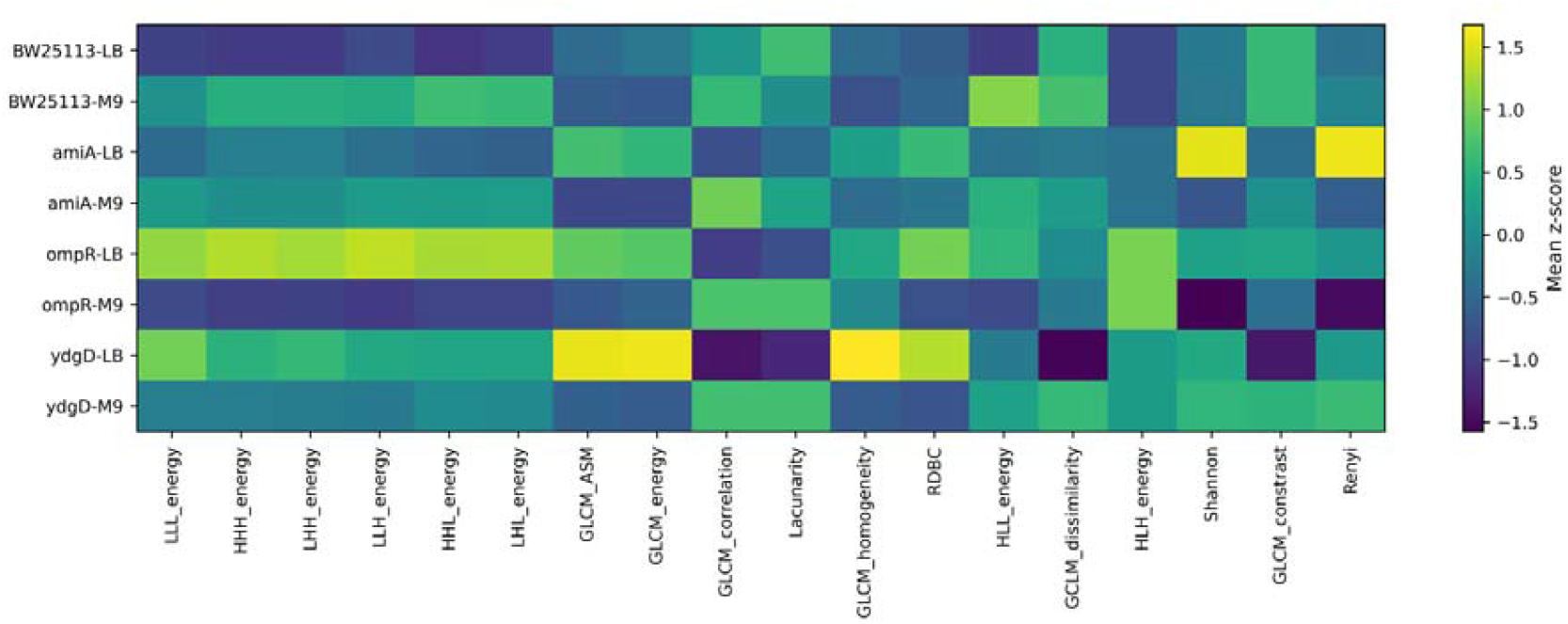
Heatmap showing mean z-scored feature values for each strain × medium combination. Rows represent experimental groups (Parental BW25113, Δ*amiA*, Δ*ompR*, Δ*ydgD* grown in LB or M9), and columns represent structural descriptors including wavelet sub-band energy parameters, GLCM texture metrics, fractal descriptors (RDBC, lacunarity), and diversity metrics (Shannon, Rényi). The colour scale indicates mean standardised values (yellow = higher than the global mean; purple = lower than the global mean). Distinct architectural fingerprints are evident for each strain-medium condition. In LB, Δ*ompR* shows broadly elevated wavelet sub-band energy and entropy-associated features, whereas Δ*ydgD* displays increased GLCM angular second momentum and GLCM energy with reduced entropy and contrast, consistent with increased local ordering. In M9, WT exhibits elevated wavelet sub-band energy with moderate homogeneity. In contrast, Δ*ompR* demonstrates reduced wavelet sub-band energy and Shannon entropy alongside relatively increased homogeneity, while Δ*ydgD* shows elevated Shannon entropy and GLCM contrast compared with its LB profile, indicating greater structural heterogeneity under nutrient limitation.

These patterns illustrate coordinated, multiscale reorganisation of biofilm architecture influenced by both cell shape and nutrient environment. The observed shifts are not monotonic across all descriptors, indicating that structural changes do not occur as uniform increases or decreases in complexity. Instead, *E. coli* biofilms comprised of different cell shapes exhibit distinct patterns of redistribution across fractal, textural, and spectral features under identical environmental conditions.

## 8. Discussion

ABC3D is presented as a framework for defining macrocolony biofilm architecture as an integrated multiscale phenotype rather than as a collection of independent structural descriptors. By unifying fractal geometry, lacunarity, entropy, grey-level texture statistics, and 3D wavelet sub-band energy features within a single reproducible workflow, ABC3D resolves architectural organisation as a coordinated spatial system. In doing so, it moves beyond conventional measures of biomass, thickness or single-parameter complexity and instead maps biofilms within a multi-dimensional architectural space.

Parameters such as fractal dimension have previously been studied in *E. coli* biofilms (25), and texture analysis has been performed in biofilms formed by *Pseudomonas aeruginosa, Pseudomonas fluorescens, Klebsiella pneumoniae* (35). However, these measurements were made from 2D projections and therefore do not fully capture the volumetric organisation of intact biofilms.

The datasets analysed here were previously assessed using 2D RDBC analysis of maximum intensity projections to quantify intra-colony channel fractal complexity (25). That study reported high fractal complexity across phenotypes and growth conditions and identified Δ*ompR* as exhibiting reduced fractal dimension when grown in M9 relative to LB. Consistent with those findings, the present 3D analysis indicates that the largest and most pronounced architectural shifts occur between Δ*ompR* and the parental BW25113 strain across nutrient conditions. However, by analysing volumetric structure directly, ABC3D reveals that these differences are not limited to changes in fractal magnitude alone but involve redistribution across wavelet sub-band energies and textural descriptors. Because maximum intensity projections compress 3D structure into a single plane, depth-dependent architectural organisation may not have been fully resolved in the earlier analysis.

In the present study, ABC3D was applied to the same datasets for analysis in 3D. This work showed the nutrient sensitivity observed in Δ*ompR* is not simply a reduction in fractal magnitude. By decomposing structure into wavelet sub-band energies and texture metrics, ABC3D reveals redistribution of spatial energy under nutrient limitation alongside increased homogeneity and reduced entropy, demonstrating that nutrient conditions reshape architectural organisation rather than merely altering fractal dimension.

The analysis also indicates that nutrient environment is the dominant determinant of architectural state, while cell shape modulates the form of that response. Strong strain × medium interactions show that structural phenotypes are conditional rather than intrinsic. The same cell shape occupies different multiscale structural configurations depending on nutrient availability, demonstrating that biofilm morphology is informed by cell shape-environment coupling (36).

This coordinated multiscale behaviour has functional implications. Gap distribution, spatial homogeneity, and textural contrast influence diffusion dynamics, metabolic stratification and mechanical stability in biofilms across species (8,10,37). When these properties shift together rather than independently, the physical microenvironment experienced by cells is altered in a structured manner. Although antimicrobial susceptibility was not directly measured here, coordinated redistribution of spatial energy and heterogeneity provides a possible structural basis for variation in stress tolerance and treatment response. Architectural state may therefore contribute to emergent phenotypes such as antimicrobial resistance through its effects on penetration, nutrient gradients and local stress buffering (38).

Beyond clinical and ecological contexts, quantitative characterisation of architectural reorganisation may also have value in biotechnology. Symmetry breaking within biofilms can generate spatially segregated sub-populations that divide labour to co-operatively synthesise complex metabolites that are difficult or costly to produce. A framework capable of quantifying coordinated multiscale structural shifts under different industrial conditions could therefore provide practical insight into how biofilm architecture supports or hampers biomanufacturing performance (39).

ABC3D extends existing biofilm analysis platforms by integrating analyses within a transparent, script-driven multivariate framework. The combination of PCA and factorial ANOVA enables reduction of high-dimensional architectural data to biologically interpretable axes while explicitly testing cell shape-environment interactions.

Several limitations of ABC3D should be acknowledged. Segmentation relied on global thresholding and may therefore not capture subtle changes in colony structure. This is particularly acute in thick macrocolony specimens where light scattering can compromise image contrast at depth. This could be improved by applying local pixel-wise thresholding methods (40). Texture analysis was implemented slice-wise rather than as a fully volumetric operation, and only single-level wavelet sub-band decomposition was applied. Incorporation of multi-level spectral analysis and fully 3D texture computation would further refine measurements.

Taken together, these findings demonstrate that biofilm architecture is best understood as a coordinated, multiscale phenotype rather than as an aggregate of independent structural metrics. By positioning biofilms within a quantitative architectural state space, ABC3D enables systematic study of how cell shape and environment jointly determine the structural organisation of colony biofilms. This framework provides a foundation for linking environmental and genetic inputs to emergent architectural properties, and for understanding how those states may subsequently shape the functional properties of biofilms.

## 10. Author statements

### 10.1 Author contributions

Gail McConnell conceived the study, developed the analysis pipeline (ABC3D), established the methodology and validation strategy, performed all formal data analysis and visualisation, curated and processed the publicly available datasets, interpreted results, and wrote and revised the manuscript. All aspects of software design, implementation, and manuscript preparation were conducted by the author.

## 10.4 Acknowledgements

Dr Manuel Banzhaf (University of Birmingham) is thanked for the original gift of the Keio collection mutants, and Dr Beatrice Bottura (Cancer Research UK Scotland Institute) and Dr Liam Rooney (University of Glasgow) are acknowledged for generating the image datasets used to demonstrate ABC3D. Thanks are also given to Dr Katherine Baxter and Dr Liam Rooney, both University of Glasgow, for critically reading the manuscript.

