## Supplementary Information 1 for "Analysis of Biofilm Complexity in 3D (ABC3D): An open-source framework for quantitative fractal, textural, and statistical analysis of colony biofilm morphology in three dimensions"

### Supplementary Information S1

#### Mathematical Definitions of Architectural Descriptors Computed by ABC3D

This document provides complete mathematical definitions of all architectural descriptors computed by ABC3D.

#### S1. Notation and preprocessing

The volumetric image is defined by

$V(z,y,x\mathbb{) \in R}$	(1)

with dimensions $z,y,x$.

Let the segmented biofilm region of interest be

$R(z,y,x) \in\{0,1\}$	(2)

The foreground voxel set is given by

$\Omega= \{(z,y,x) : R(z,y,x)=1\}$	(3)

with a total voxel count of

$N = zyx$.	(4)

#### S2. Intensity Statistics

The mean intensity is given by

$\mu= (1/|\Omega|) \Sigma V(r)$	(5)

With the standard deviation given by

$\sigma= \sqrt{(1/|\Omega|) \Sigma{(V(r) - \mu)}^{2}}$	(6)

#### S3. Entropies

Shannon and Rényi entropies are given by

$H_{Shannon}=-\sum_{i} p_{i}\log p_{i}$	(7)

and

$H_{Rényi,q=2}=-log\sum_{i} p_{i}^{2}$	(8)

respectively.

#### S4. Box-Counting Dimension

#### For dyadic box size s, $\boldsymbol{N(s)}$ is the number of boxes containing a minimum of one foreground voxel thus

$ln N(s) = D ln(1/s) + c$	(9)

$R^{2} = 1 - [\Sigma{(y_{k} - ŷ_{k})}^{2} / \Sigma{(y_{k} - ȳ)}^{2}]$.	(10)

#### S5. Lacunarity

For dyadic box size s, lacunarity is given by

$\Lambda(s) = Var[M(s)] / ({Mean[M(s)]}^{2}) + 1$	(11)

and the lacunarity slope from regression is given by

$ln \Lambda(s) = m ln s + b$	(12)

#### S6. Fourier Energy

The Power spectrum is given by

$P(k) = {|F(k)|}^{2}$.	(13)

The total energy is given by

$E_{total} = \Sigma P(k)$.	(14)

The high frequency ratio is

$R_{HF} = (E_{total} - E_{low}) / E_{total}$	(15)

#### S7. GLCM Metrics

After quantisation to L grey levels, the normalized co-occurrence matrix $P\left( i, j \right)$ can be computed.

$Angular second moment \left( ASM \right)=\sum_{i,j} P\left( i,j \right)^{2}$	(16)

$Contrast=\sum_{i,j} \left( i-j \right)^{2}P\left( i,j \right)$	(17)

$Correlation=\frac{\sum_{i,j} \left( i-\mu_{i} \right)\left( j-\mu_{j} \right)P\left( i,j \right)}{\sigma_{i}\sigma_{j}}$	(18)

$Dissimilarity=\sum_{i,j} \left| i-j \right|P\left( i,j \right)$	(19)

$Homogeneity=\sum_{i,j} \frac{P\left( i,j \right)}{1+\left( i-j \right)^{2}}$	(20)

#### S8. Wavelet Sub-band Energies

Prior to transformation, the intensity volume is mean-centred within the segmented region of interest (ROI). The mean intensity is given by Equation (5).

The wavelet transform is applied to the ROI-masked, mean-centred signal

$X\left( r \right)=\left\{ \begin{matrix} (V\left( r \right)-\mu, R\left( r \right)=1 \\ 0, R\left( r \right)=0 \end{matrix} \right.$	.	(21)

A single-level 3D discrete wavelet transform using the Daubechies db2 wavelet with symmetric boundary extension is performed

$\left\{ C_{abc} \right\}=W_{db2}\left\{ X \right\},$	(22)

where

$a, b, c \in\left\{ L, H \right\}$	(23)

denote low-pass $(L)$ and high pass $(H)$ filtering along the $z$, $y$, and $x$ axes respectively, giving the 8 sub-bands $LLL, LLH, LHL, LHH, HLL, HLH, HHL,$ and $HHH$.

The energy of each sub-band is defined as

$E_{abc}= \sum_{i, j, k} C_{abc}\left( i, j, k \right)^{2}$	(24).
